# Nanoparticle Metal Mass Uptake Governs Radiosensitizing Efficacy Across 2D, 3D, and In Vivo Models

**DOI:** 10.1101/2025.08.13.670090

**Authors:** Lukas R.H. Gerken, Laurin G.S. Schaller, Rüveyda Dok, Selina Camenisch, Alexander Gogos, Sebastian Habermann, Sandra Nuyts, Inge K. Herrmann

## Abstract

Despite extensive efforts to develop nanoparticle-based radioenhancers, clinical translation remains limited, partly due to the lack of physiologically relevant *in vitro* models. To address this gap, we developed a 3D spheroid model of head and neck cancer using FaDu cells and compared it directly to a corresponding *in vivo* model in a radiotherapy setting. The spheroids exhibited key tumor-like features, including the formation of a hypoxic core and growth kinetics comparable to *in vivo* tumors. Importantly, the model allowed for long-term monitoring of tumor growth and radiation response. Upon X-ray irradiation, dose–response behavior in spheroids mirrored that observed *in vivo*. Furthermore, TiO₂, HfO₂, and Au nanoparticles demonstrated consistent radiosensitization effects in both systems when matched for uptake mass. In contrast, conventional 2D clonogenic assays failed to predict *in vivo* performance, likely due to their lower radioresistance and unrealistic nanoparticle exposure conditions. This study introduces a robust, scalable, and clinically compatible 3D *in vitro* platform for preclinical screening of nanoparticle radioenhancers. The system may offer streamlining of development pipelines and support the 3R principles of reduction, replacement, and refinement in radiation oncology research.

## Introduction

Radiation therapy is a cornerstone in the multimodal treatment concept of cancer with approximately 50% utilization rate in all cancer cases and a projected increase in global clinical need by 2035.^1^ Radiotherapy (RT) has seen continuous technological and treatment-planning innovation for decades, however, normal tissue dose limits X-ray based RT from achieving higher efficacy.^2,3^ To further improve the efficacy of X-ray RT, radioenhancers and radiosensitizers that amplify the deposited dose locally or sensitize cancer cells to ionizing radiation are investigated.^4^

While a plethora of radiosensitizer nanoparticles have been investigated preclinically with great success, translation of these candidates is extremely slow and only a few materials are currently investigated in clinical trials, such as NBTXR3 (HfO_2_) or Gd-AGUIX.^4–9^ One of the bottlenecks of the translation of *in vitro* radioenhancer nanomaterials is the use of 2D cell culture settings, such as the gold standard clonogenic assay.^10,11^ Such two-dimensional cultures allow high-throughput screening, but are still time-consuming and lack physiological relevance including complex signals, structures, nutrient and oxygen gradients.^12^ To translate findings into clinics animal experiments are therefore necessary, which are costly, pose ethical strains, and often do not reflect the same efficacy as found in 2D cell culture systems. To overcome these shortcomings, 3D cell culture systems from simple single cell to multicellular spheroids or organoids made from cell lines or primary cells with increasing complexity have been proposed.^13,14^ Such methods can also be used for the high-throughput assessment of nanoparticle radioenhancement properties, allowing a rational and data-driven material development.^15^ Since 3D cell cultures show much more resistant dose-response relationships compared to 2D cultures,^16^ they seem to be more realistic and comparable to *in vivo* scenarios.

Amongst the various types of cancers, head and neck cancer (HNC) are of particular interest due to their increasing incidence rates as well as their anatomical complexity.^17,18^ Radiotherapy plays a crucial role in the treatment of HNC and is used alone, or in combination with surgery, chemo- or immunotherapy.^18–20^ Despite the multimodal therapy efforts, the 3-year local recurrence rate is as high as 50%.^20^ Problematic limitations of RT success in HNC are treatment related toxicity effects and radioresistance due to high prevalence of hypoxia.^21–23^ Therefore, new treatment options that can increase the efficacies of RT in HNC are necessary.^24,25^ In this study, we investigated the predictive value of using 3D spheroids for nanoparticle radioenhancement therapy in HNC. We first characterized the cell uptake and radiosensitization effects of TiO_2_, HfO_2_ and Au nanoparticles in FaDu cells in a 2D culture setting using the clonogenic assay. Thereafter, we characterized FaDu spheroids in terms of physiological characteristics, growth and dose-response with and without nanoparticles. Finally, we quantified the radiosensitization effects of TiO₂, HfO₂, and Au nanoparticles at comparable levels of cellular uptake across 2D cultures, 3D spheroids, and FaDu tumor xenografts, along with analysis of the nanoparticle biodistribution. Collectively, our findings demonstrate that 3D spheroids reliably recapitulate *in vivo* radiosensitization responses, supporting their use as predictive preclinical models. Moreover, we identify nanoparticle metal mass uptake as a key determinant of radiosensitization efficacy across all experimental systems.

## Results and Discussion

### Nanoparticles, cell uptake and cell monolayer radiosensitization

All nanoparticles used in this study, including commercially available citrate-stabilized 50 nm Au nanoparticles, and flame spray pyrolysis (FSP)-synthesized HfO_2_ and TiO_2_ nanoparticles (∼5–10 nm primary particle size, forming 10–100 nm agglomerates) were characterized by Transmission electron microscopy (TEM) (Figure 1a).

**Figure 1:**
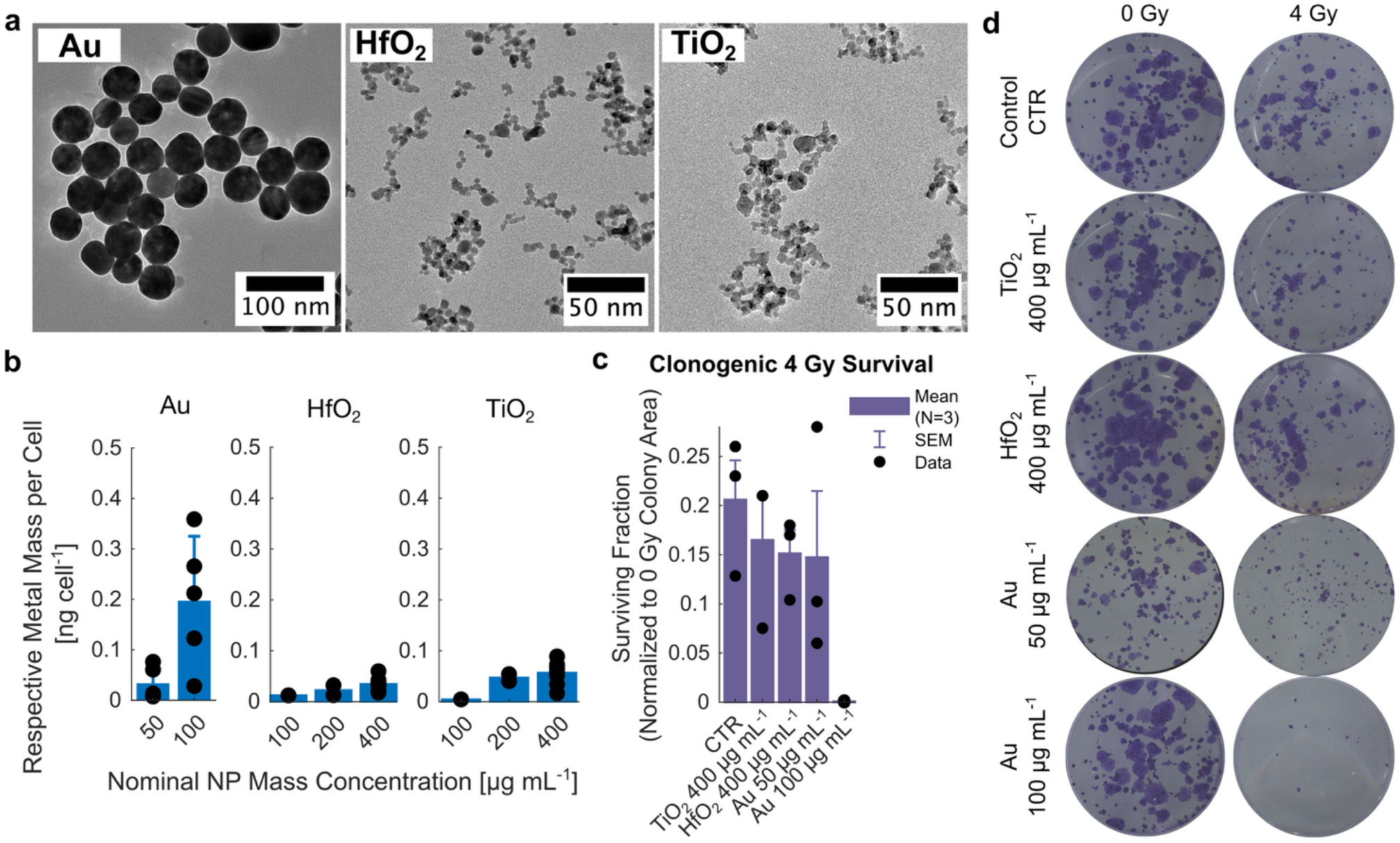
TEM images of Au, HfO_2_ and TiO_2_ nanoparticles (a); Cellular metal mass uptake after 24h nanoparticle incubation of FaDu cell monolayers (b); Clonogenic cell survival of Fadu cells treated with nanoparticles and 4 Gy 150 kVp X-ray irradiation (c), data shown as Mean ± SEM from N = 3 different biological experiments; Exemplary image scans of colonies post 0 or 4 Gy irradiation (d).

To assess nanoparticle-mediated radioenhancement in the simplest possible system, we performed 2D cell culture experiments using the FaDu cell line, derived from a human pharyngeal squamous cell carcinoma. After 24 hours of incubation with varying nanoparticle concentrations, Au nanoparticles exhibited significantly higher cellular uptake than the metal oxides (Figure 1b). Quantitative analysis of intracellular metal content by ICP-OES revealed that exposure to 50 µg mL⁻¹ Au or 400 µg mL⁻¹ of TiO_2_ or HfO_2_ nanoparticles resulted in a comparable uptake of approximately 0.5 ng per cell. Doubling the Au concentration to 100 µg mL⁻¹ in turn led to a fourfold higher uptake. As shown in our previous studies, both FSP-synthesized metal oxides and Au nanoparticles are internalized by epithelial cancer cells via endocytic pathways and accumulate in ∼500 nm-sized intracytosolic endosomal agglomerates.^26,27^ Such intracellular nanoparticle uptake has been shown to be size dependent, with an uptake maximum for nanoparticles with around 50 nm diameter,^28–32^ which has been explained by favorable cell membrane adhesion properties and optimal energetic conditions of nanoparticle wrapping.^33–35^

Nanoparticle incubation in 2D FaDu cell cultures resulted in weak radiosensitization, except for the highest Au nanoparticle concentration (100 µg mL⁻¹), which significantly reduced survival after 4 Gy irradiation due to ∼4× higher cellular uptake and the strong dose-enhancing properties of gold. Cell survival was assessed by quantifying colony-covered well area, a robust metric for difficult-to-count colonies.^36,37^ Approximately 20% of control cells survived a 4 Gy irradiation. At comparable metal mass uptake (∼0.5 ng cell^−1^), the radiosensitization effect resulting in ∼15% cell survival was similar across nanoparticle types. The same trend has been confirmed with a viability assay 10 days post irradiation (ESI, Figure S1). Radiosensitization effects correlated with metal uptake, particularly for Au nanoparticles, and were similar across materials at comparable metal mass. Our results align with previous reports on HfO_2_ (NBTXR3) nanoparticles, for which dose enhancement ratios between 1.1 – 2.5 have been documented in FaDu cells with a 200 kVp-X-ray dose of 4 Gy,^38^ though observed at lower nominal nanoparticle concentrations (100 – 800 µM, 20 – 170 µg mL^−1^), likely due to higher uptake of 50 nm sized nanoparticles. Higher radiation doses did not lead to higher sensitization effects relative to control cells treated with the same X-ray dose (ESI, Figure S2).^38^ To better reproduce tissue level properties, we established and characterized 3D FaDu spheroids to better model realistic nanoparticle uptake and radioenhancement responses.

### Spheroid characterization and response to radiotherapy

To characterize the 3D phenotype of FaDu cells during spheroid growth, we performed live/dead and hypoxia staining (Hoechst, Propidium Iodide, Hypoxia Green) at days 3 and 10 post-seeding. By day 3, spheroids (∼400 µm diameter) showed a hypoxic but viable core with minimal cell death and no necrotic center (Figure 2a). By day 10 (∼600 µm), a necrotic core (<300 µm) had formed, and only the outer 30–40 µm cell layer remained normoxic. This aligns with literature reporting necrosis at spheroid diameters ≥500 µm due to limited oxygen diffusion (>150 µm from the surface) and high metabolic demand of cancer cells.^39,40^ Hypoxia Green staining confirmed that only peripheral cells experienced oxygen levels >5%. These features highlight the physiological relevance of the spheroid model for solid tumor studies, distinguishing it from 2D monolayers.

**Figure 2:**
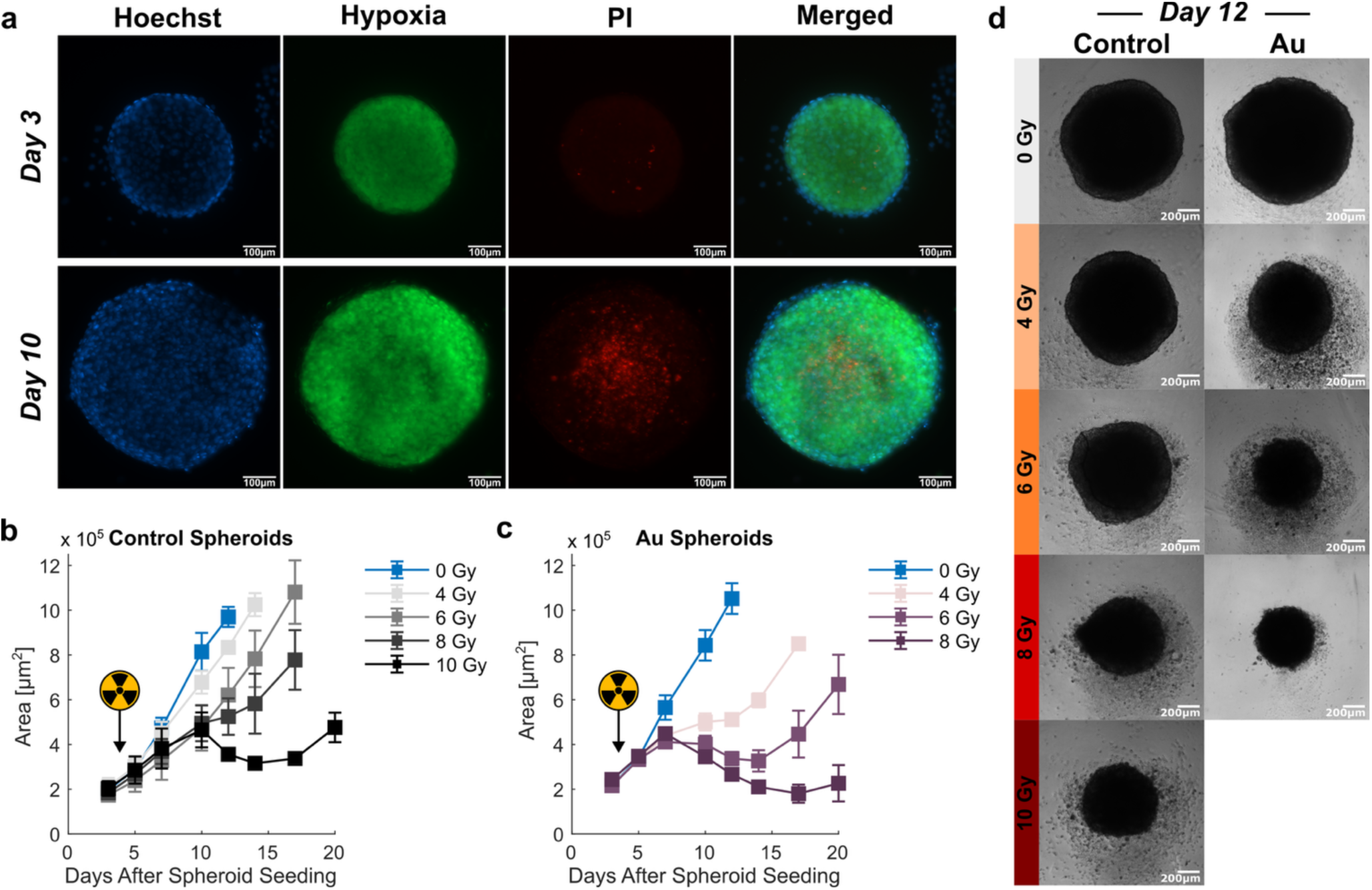
Spheroid characterization on day 3 and day 10 after spheroid seeding using Hoechst (viable cells, blue) Hypoxia (green) and Propidium Iodine (PI, dead cells, red) stainings (a); Spheroid area growth curves of control (c) and Au treated (c) spheroids irradiated on day 4 after spheroid seeding with 0, 4, 6, 8 or 10 Gy 150 kVp X-rays (n=4); Bright field images of control and Au-treated spheroids irradiated with 0, 4, 6, 8 or 10 Gy imaged on day 12 after spheroid seeding (d).

To assess radiosensitivity, spheroids were irradiated on day 4 with 0–10 Gy X-rays (150 kVp). Growth delay was dose-dependent (Figure 2b): after 4 Gy, delay became apparent on day 10, while 6, 8 and 10 Gy caused earlier and more pronounced growth delay effects (see ESI Table S1). Four-fold area increases (∼8000 µm^2^) occurred on days 10 (0 Gy), 12 (4 Gy), 14 (6 Gy), and 17 (8 Gy). Compared to 2D cultures, spheroids showed greater radioresistance (in line with the literature^41^): 4 Gy reduced 2D cell survival by 80%, but only ∼20% growth inhibition was seen in 3D spheroids by day 12.

Following the spheroid model establishment, the FaDu spheroid model was utilized to assess the robustness of nanoparticle RT enhancement efficacy in nanoparticle treated spheroids (Figure 2c). For this, Au nanoparticle treatment was chosen as a robust and gold-standard benchmark. Non-irradiated control and Au-treated spheroids had ∼2000 µm^2^ initial growth area 3 days post spheroid seeding and revealed similar growth behavior for nanoparticle exposed and untreated spheroids, indicating the absence of relevant nanoparticle toxicity. In contrast Au nanoparticle treatment in combination with RT lead to a severe delay of spheroid growth compared to control spheroids, indicating radiation dose enhancing effects. In fact, very significant growth delay was already found on day 7 for ≥ 4 Gy X-ray treatments. Four-fold spheroid growth area (∼8000 µm^2^) of Au-treated spheroids was reached approximately on days 10, 17 and 20 when treated with 0, 4 and 6 Gy, respectively. Therefore, a 4 Gy treatment of Au-containing spheroids showed a treatment outcome comparable to an 8 Gy treatment of control spheroids. The bright field images on day 12 post spheroid seeding demonstrate that 0 Gy treated control and Au spheroids had a similar size, while the 4 and 6 Gy treated Au spheroids had comparable sizes to the 8 and 10 Gy treated control spheroids, respectively (Figure 2d). Therefore, the nanoparticle dose enhancement effect was relatively robust across X-ray doses and was approximately two-fold with a 4 or 6 Gy X-ray treatment.

### Nanoparticle cell uptake and radiosensitization in cell spheroids

Based on the findings of Au-nanoparticle radioenhancement in spheroids, the following protocol was defined allowing to investigate radioenhancing properties of several nanoparticles in the spheroid model with one X-ray dose (Figure 3a). The standardized experimental design of the following nanoparticle radioenhancement study in spheroids included a 24h nanoparticle loading in 2D cell monolayers, a 3-day spheroid formation phase, 6 Gy irradiation treatment, nanomaterial uptake quantification, growth analysis and viability assessments.

**Figure 3:**
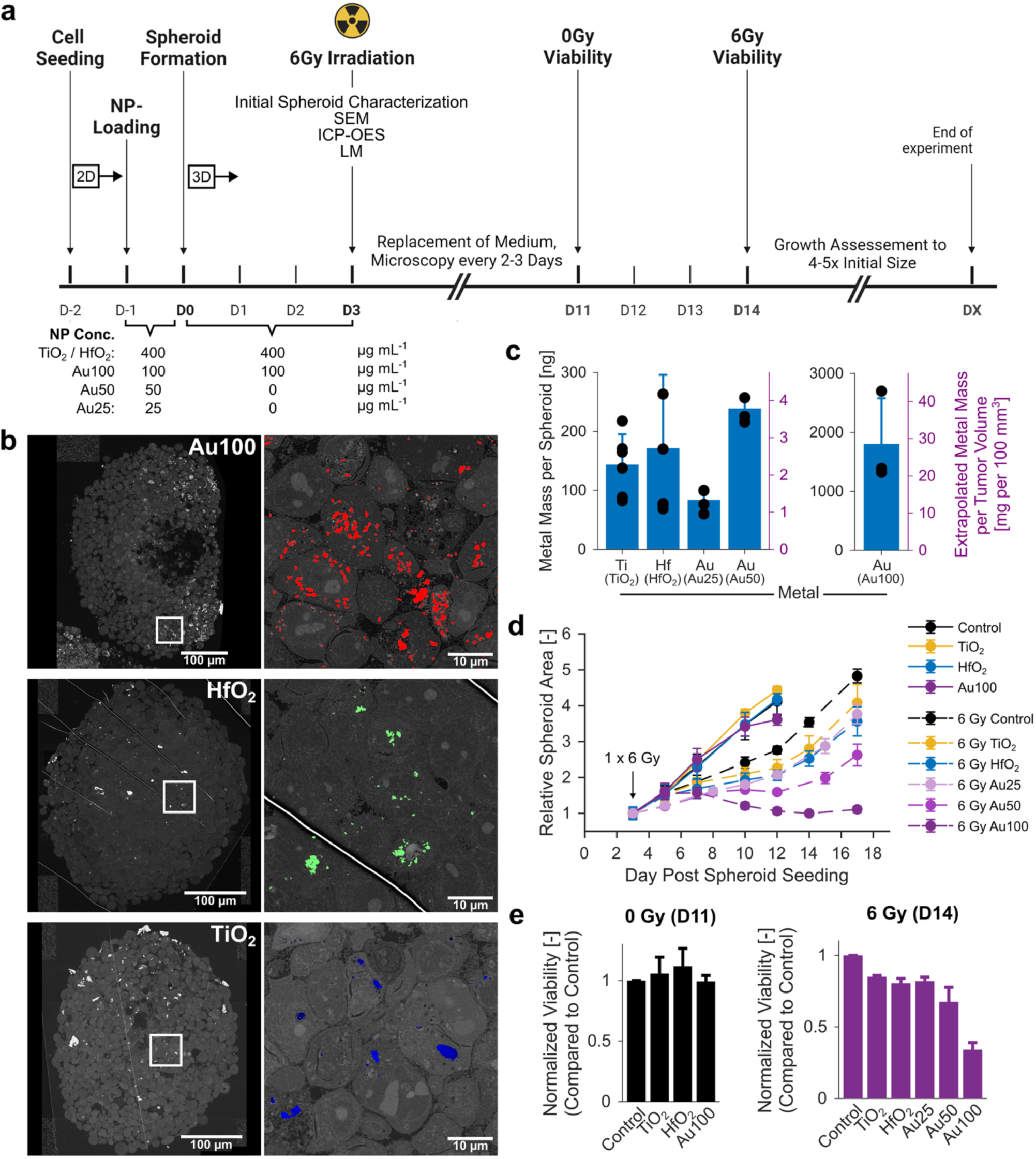
Schematic illustration of spheroid irradiation experiments with indications of relevant concentrations and timepoints of nanoparticle and irradiation treatment, medium exchanges, and viability and growth curve assessments, D: Day Post Spheroid Seeding (a); Nanoparticle distribution in spheroids visualized with SEM-imaging at D3; insets (right column) show magnified area with overlayed, colored segmentation of nanoparticles (Au: red; HfO_2_: green, TiO_2_: blue) (b); Metal mass per spheroid as determined by ICP-OES on D3 (Mean ± SD, N=3) (c); Spheroid area growth curve measured over 17 days (D3 – D17) expressed as relative spheroid area (RSA) (d); Viability of non-irradiated control and nanoparticle-treated spheroids on day 11 (D11) and of irradiated spheroids on day 14 (D14) as assessed by a ATP-based assay (Mean ± SEM, N=3) (e).

Scanning electron micrographs taken on the day of irradiation revealed a heterogeneous distribution of nanoparticles within the spheroids (Figure 3b). Uptake varied significantly between individual cells: while some exhibited minimal or no internalization, others showed substantial accumulation of aggregated nanoparticles. Clusters were also observed in the intercellular spaces and on the membranes of peripheral cells, with aggregate sizes spanning a broad range. Metal uptake per spheroid was quantified using Inductively Coupled Plasma Optical Emission Spectroscopy (ICP-OES). For TiO_2_, HfO_2_, Au50, and Au25-treated spheroids, uptake ranged between approximately 80 and 240 ng of metal per spheroid. This corresponds to an extrapolated concentration of 1–4 mg per 100 mm^3^ tumor volume. According to Nanobiotix’s recommendations and previous studies on intratumoral radioenhancer injections, around 25–33% of a 54.2 mg mL^−1^ nanoparticle solution is typically administered, equating to roughly 1.3–1.8 mg per 100 mm^3^ tumor volume.^42,43^ Thus, the observed uptake in TiO_2_, HfO_2_, Au25, and Au50-treated spheroids falls within a realistic and translatable range for *in vivo* application. In contrast, Au100-treated spheroids exhibited an unrealistically high uptake of 20–30 mg per 100 mm^3^. For Au100, TiO_2_, and HfO_2_, nanoparticles were also present in the medium during spheroid formation, underscoring the preferential uptake of 50 nm gold particles even during the early aggregation phase. These conditions also led to greater variability in the uptake data, as reflected by the higher standard deviations compared to Au25 and Au50 treatments. This variability likely results from nanoparticle aggregation on the spheroid surface during early formation, potentially leading to an overestimation of internal uptake, as confirmed by optical microscopy. High uptake in spheroids treated with 100 µg mL^−1^ Au nanoparticles was expected, as previous studies using 50 µg mL^−1^ Au also reported milligram-range accumulation per spheroid.^15^ Given that nanoparticle uptake levels were within realistic *in vivo* ranges, we proceeded with RT on nanoparticle-loaded spheroids and monitored growth over time. Nanoparticles alone had no significant effect on spheroid growth compared to controls. However, combining TiO_2_, HfO_2_, or Au25 with 6 Gy irradiation led to delayed spheroid growth relative to irradiated controls. Uptake-dependent enhancement was evident for all Au treatments, with Au100 almost completely suppressing growth for 14 days post-irradiation. For comparable metal masses, growth delay was similar regardless of the metal type. To assess whether reduced growth reflected lower cell viability, a 3D metabolic assay was conducted. On day 11, non-irradiated spheroids showed similar viability across all groups. In irradiated spheroids, viability dropped by ∼15–20% for HfO_2_-, TiO_2_-, and Au25-treated samples, and by ∼40% and ∼70% for Au50 and Au100, respectively, compared to irradiated control spheroids. These viability trends correlated well with growth curves, confirming that size reduction observed via microscopy was mirrored by decreased metabolic activity.

### Nanoparticle Distribution in vivo

Following the in-depth characterization of nanoparticle enhancement effects in spheroids, we conducted an *in vivo* study using intratumoral injections, analogous to previous protocols with CE-approved NBTXR3 nanoparticles for optimal comparability.^42,43^ FaDu cells were subcutaneously injected into both flanks of nude mice, and ∼1 mg of nanoparticles per 100 mm³ tumor volume was administered into both tumors.

To assess nanoparticle biodistribution, we employed SEM, H&E staining, and cone beam CT. For successful delivery, nanoparticles were stabilized using BSA^44^ (ESI, Figure S2). Correlative SEM and histology on day 1 post-injection revealed widespread but heterogeneous nanoparticle distribution, including agglomerates (Figure 4a). In the case of Au nanoparticles, both agglomerates and individual particles were detectable via SEM and histology, often adjacent to intact nuclei. Due to their strong optical properties, Au nanoparticles were also clearly visible in histological sections. CT imaging on days 1–5 confirmed the presence and retention of nanoparticles within tumors (Figure 4b, ESI Figures S3–S5). HfO_2_ and Au provided strong contrast, while TiO_2_ gave weaker but still distinguishable signals. Due to lower resolution, CT primarily visualized regions with high nanoparticle concentration. Nonetheless, signal persistence over five days suggests prolonged tumor retention. Metal quantification in organs and tumors (6–24 h post-injection) confirmed most of the metal was retained in the tumors, with similar levels across nanoparticle types (Figure 4c). Some accumulation was detected in the liver, indicating hepatic clearance as a likely elimination pathway. Lymphatic drainage followed by renal (for ≤5 nm) or hepatobiliary excretion are known nanoparticle clearance routes.^45,46^ Literature confirms high intratumoral retention post-injection, with variability depending on tumor type. For example, Zhang et al. demonstrated long-term retention of NBTXR3 across multiple tumor models via µCT.^43^ Similarly, a phase I trial in head and neck cancer reported NBTXR3 retention over seven weeks without leakage.^47^ Minimal systemic distribution and lack of renal clearance have been documented in both preclinical and clinical studies.^47–49^ Our findings confirm strong retention of intratumorally injected nanoparticles, with minimal off-target accumulation and extended residence times in tumor tissue.

**Figure 4:**
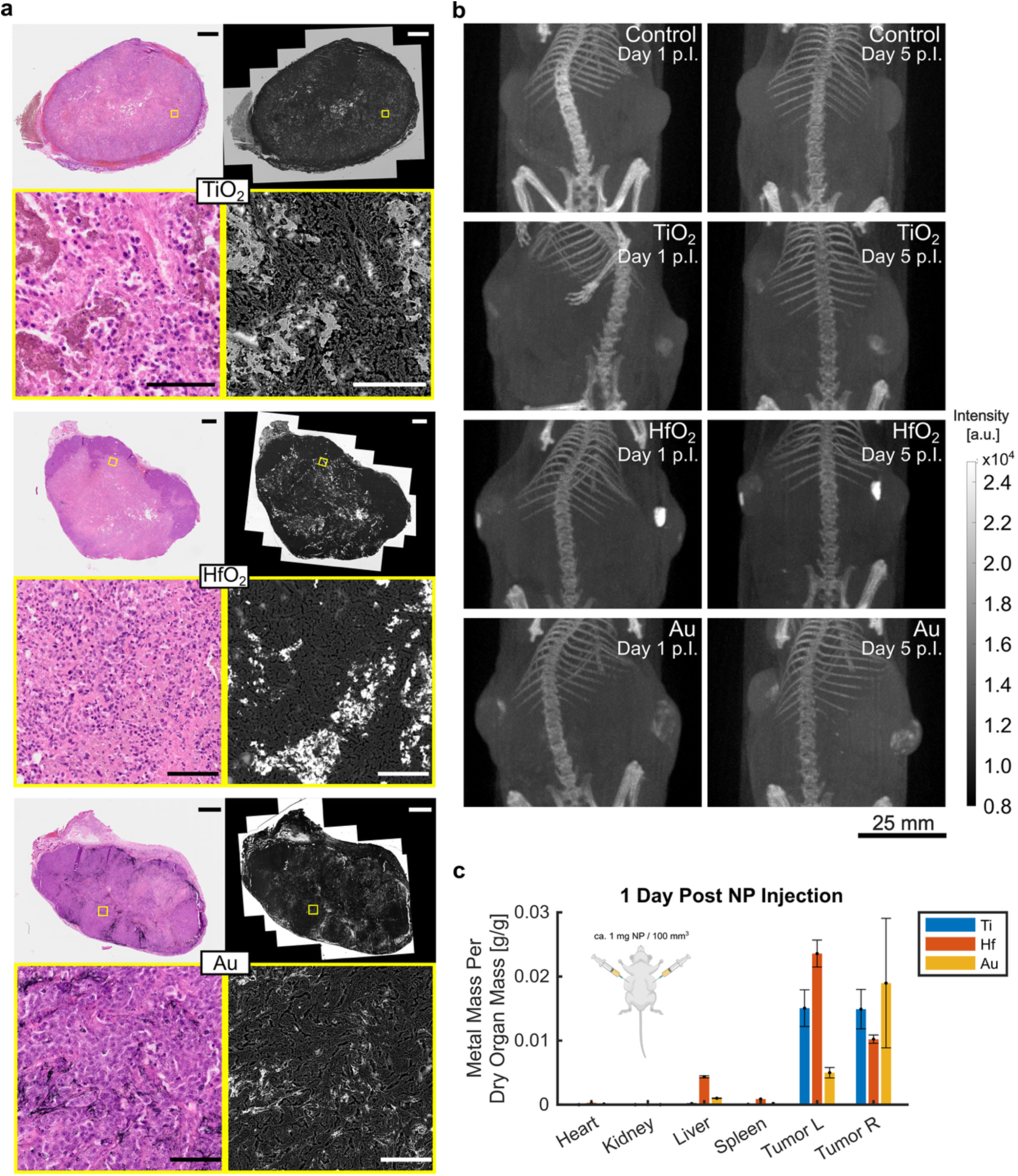
LM and SEM overview and zoom images showing nanoparticle distribution throughout a tumor (FaDu) slice correlated to the same H&E stained histology image (tumor harvested 1 day (HfO_2_, TiO_2_) or 6 hours (Au) post nanoparticle injection) – Scale Bars: 1000 µm in overview images, 100 µm in zoomed images (a); Maximum intensity projection of cone beam CT images of day 1 (start of radiotherapy) and day 5 (end of radiotherapy) post intratumoral nanoparticle injection (p.I.: post Injection) (b); Recovered metal mass per dried organ mass of mice injected with nanoparticles and sacrificed 1 day (HfO_2_, TiO_2_) or 6 hours (Au) post nanoparticle injection (n = 1 mouse per nanoparticle, L: Left, R: Right) (c).

### Nanoparticle Radiosensitization in vivo

To confirm the ability of the various nanoparticles to amplify RT treatment *in vivo*, we performed 5 x 2 Gy RT during day 1 – 5 post nanoparticle injection (Figure 5). In the absence of RT, the injection of nanoparticles did not affect tumor growth, confirming the non-toxic nature of these nanoparticles that has also been observed *in vitro* in spheroids (Figure 5a). RT treatment led to a marked tumor growth delay starting approximately on day 8 post nanoparticle injection, or on day 3 post the last RT dose. This observation is congruent with the tumor growth delay behavior of *in vitro* spheroids. Nanoparticle treated tumors overall showed better tumor control and improved survival compared to RT alone. All nanoparticles increased the median survival from 30 days (RT) to 34 days (RT+NPs) (Figure 5b-c). Furthermore, the retained metal mass per tumor at the end of the experiment (34 days post NP injection) was found at comparable levels, around 0.3 – 0.5 wt% (Figure 5d), suggesting that tumor growth control improvement *in vivo* with kVp-X-ray treatment until day 34 is governed by the intratumoral total metal mass rather than the type of metal. Nevertheless, the underlying enhancement mechanisms might be different for each nanoparticle and explained by physical dose enhancement with high-Z Au and HfO_2_ nanoparticles, and by catalytic chemical radiosensitization with TiO_2_ nanoparticles.^50,51^

**Figure 5:**
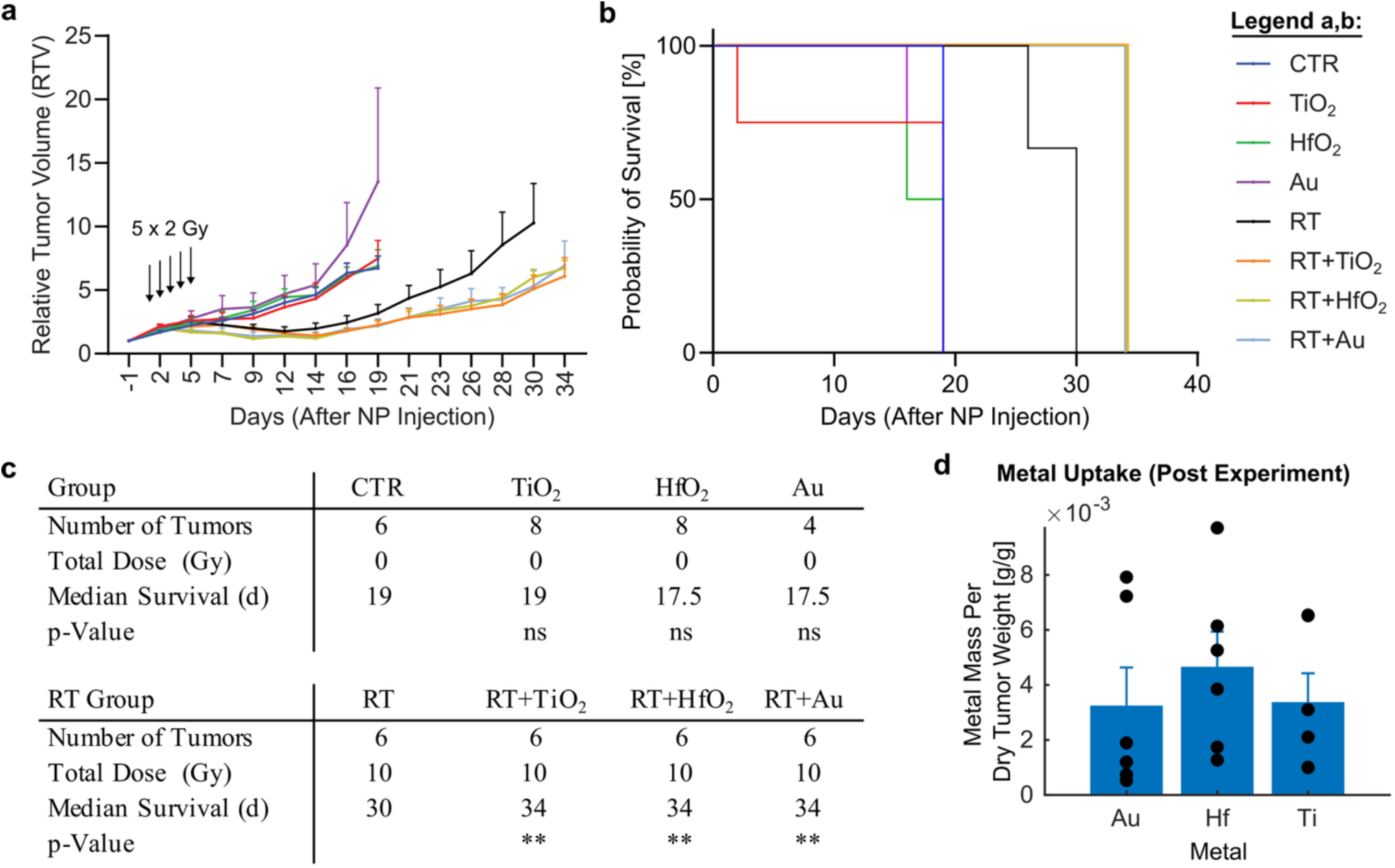
Tumor growth curves (a) and survival probability (b) of FaDu tumors (Mean ± SEM, n = 6). X-axis expressed in days after nanoparticle (NP) injection; radiotherapy (RT, 5 x 2 Gy) was performed on days 1 to 5 post NP injection; Statistical results of survival probability (c); CTR: Control; d: days; ns: not significant (p > 0.05); **: 0.001 ≤ p < 0.01. Recovered metal mass per dry tumor weight in the RT+NP groups after sacrifice of mice (day 34) (d).

The growth delay results observed in this study align with those reported by Zhang et al. for NBTXR3+RT in the FaDu model, where increased overall survival was demonstrated.^43^ However, their limited observation period (<20 days) precluded detection of statistically significant treatment effects, in contrast to our study, which revealed significant growth delays for both metal oxide and gold nanoparticles. Notably, we directly compared multiple nanoparticle types and found comparable radiosensitization effects at similar *in vivo* and *in vitro* uptake levels. While single-dose *in vitro* irradiation is not directly representative of clinical radiotherapy, which typically employs fractionated dosing (e.g., 5 × 2 Gy),^52^ we observed analogous trends. *In vivo*, RT (5 × 2 Gy) induced a tumor growth delay (Δt) of ∼9 days to reach 5×RTV (relative tumor volume) compared to controls (day 14–23, Figure 5a). This was similar to the ∼7-day delay observed in spheroids treated with a single 8 Gy dose, achieving 3×RSA (Relative Spheroid Area, RSA = ∛(RTV²); day 8–15, Figure 2b). However, spheroids grew ∼3× faster than *in vivo* tumors (control Δt(5×RTV) ≈ 15 days (Fig. 5a) vs. control Δt(3×RSA) ≈ 5 days, Fig. 3d). Thus, the 1 × 6 Gy spheroid model approximated the tumor growth kinetics of the 5 × 2 Gy *in vivo* protocol reasonably well. Finally, combining nanoparticles with RT resulted in growth delays of ∼7 days *in vivo* (Figure 5a, day 23–30) and ∼2–3 days in spheroids (Figure 3d, day 12/13–15) compared to RT alone, with comparable nanoparticle uptake per tumor/spheroid volume.

## Conclusion

Driven by the lack of data on the nanoparticle radioenhancement effects in 3D cell culture and the limited clinical translation despite extensive material screening, we developed a spheroid model of head and neck cancer using FaDu cells and compared it to the corresponding *in vivo* model. Model characterization revealed the formation of a hypoxic core, making it a suitable system for studying solid, hypoxic tumors. The spheroids showed growth kinetics similar to *in vivo* tumors and enabled long-term monitoring of size changes. X-ray dose–response relationships in spheroids closely matched those observed *in vivo*. TiO_2_, HfO_2_, and Au nanoparticles produced comparable radiosensitization effects at similar uptake levels in both models. In contrast, 2D clonogenic assays underestimated *in vivo* responses due to lower radioresistance and unrealistic uptake conditions. Overall, we present scalable spheroid-based *in vitro* platform for evaluating nanoparticle-based radioenhancers prior to *in vivo* validation of selected candidates. It is compatible with clinical treatment machines (e.g., 6 MV Linac) and supports high-throughput testing, helping to accelerate clinical translation while reducing the need for animal experiments in line with the 3R principles of reduction, replacement, and refinement.^53^

## Materials and Methods

### Nanomaterials and Characterization

All nanoparticles used in this study were imaged using transmission electron microscopy (TEM) confirming previous characterizations and are depicted in Figure 1a. Spherically shaped gold (Au) nanoparticles were 50 nm sized and commercially available as aqueous suspension stabilized with citrate molecules (BioPure, nanoComposix Inc., USA). Therefore, they were readily suspensible in/miscible with cell medium easing their use in cell culture experiments. Hafnium dioxide (HfO_2_) nanoparticles were prepared by Avantama AG via an industrially scalable Flame Spray Pyrolysis (FSP) synthesis method.^26^ The primary particle size was around 5 – 10 nm in diameter, as determined by XRD, BET and TEM, and nanoparticles occurred in fractal agglomerates of 10 – 100 nm (DLS) which is a well-studied feature of the particular synthesis method.^26^ Titanium dioxide (TiO_2_) nanoparticles were very comparable in shape, size and occurrence to HfO_2_ nanoparticles and were also synthesized by the FSP method.^26^ To disperse FSP-made metal oxides in cell medium to study cellular interactions, bath sonication of the aqueous nanoparticle solution was required beforehand.

### Cell Line and Cell Handling

The FaDu (HTB-43) cell line, a human squamous cell carcinoma cell line derived from the epithelial tissue of a hypopharyngeal tumor was used in this study. It was obtained from the American Type Culture Collection (ATCC, USA). Cells were cultured in cell growth medium consisting of Minimum Essential Medium Eagle (Sigma-Aldrich, Switzerland) supplemented with 10% FCS (Sigma-Aldrich, Switzerland), 1% Penicillin-Streptomycin (Sigma-Aldrich, Switzerland), 1% L-glutamine (Sigma-Aldrich, Switzerland), 1% non-essential amino acids (PAN Biotech, Germany) and 1% sodium pyruvate (Sigma-Aldrich, Switzerland). FaDu cells were maintained in a humidified incubator at 37°C, 5% CO_2_ and cultured to 80%-90% confluence before being split. For experiments, cells between passages 5 and 30 were used.

### Nanoparticle treatment of Cell Monolayers

For each experimental condition, 1 - 1.5 × 10^5^ cells were seeded in 2.7 mL growth medium into 6-well plates and allowed to adhere for 24 hours. The following day, 10x concentrated nanoparticle dispersions were freshly prepared from Au nanoparticle suspensions, or from TiO_2_ or HfO_2_ powder that were weighted and suspended in milipore water followed by bath sonication. Then, 0.3 mL of the concentrated NP dispersions were added to the 2.7 mL cell medium to reach the desired NP concentrations as indicated in the main text. Controls were treated with 0.3 mL milipore water.

### Cell Monolayer Irradiation, Viability and Clonogenic Assay

After 24 h nanoparticle or control treatment of FaDu cells, 6 well plates were placed in a PMMA phantom described elsewhere (2 slabs with 4 cm thickness)^50^ and irradiated with 150 kVp X-rays (Seifert ISOVOLT 450, GE Sensing & Inspection Technologies GmbH, Germany) at a dose rate of ∼1.5 Gy min^−1^. 2 h after irradiation, cells were washed twice with PBS and detached using 450 µL 0.5% Trypsin-EDTA (Sigma Aldrich, Switzerland). After 5 min, trypsination was stopped and cells were transferred to tubes using 2 x 500 µL growth medium. Cells were palleted using centrifugation (200g, 5 min), redispersed in 450 µL growth medium, counted, and seeded in their respective amounts (0 Gy: 300 cells, 2 Gy: 600 cells, 4 Gy: 1200 cells, 6 Gy: 3333 cells) in duplicates in 6 well plates with a total growth medium volume of 3 mL per well. After 2 weeks, cells were fixed and stained using 0.5 mL of 5% glutaraldehyde and 0.5 mL 0.4% crystal violet. For cell viability assessment after cell counting, 500 cells were seeded into 48 well plates in quadruplicates and viability was assessed after 10 days using the CellTiter-Glo® assay (Promega, Switzerland) and a self-developed PEEK light shielding 48-well plate adapter protecting from well-to-well cross talk during luminescence reading.^50,51^

### Spheroid Formation, Nanoparticle Treatment and Irradiation

The day prior to spheroid formation cell monolayers were loaded with nanoparticles or control vehicle as described before. After 24 hours NP loading, nanoparticle containing cells were seeded in ultra-low attachment plates to let them form spheroids to progress into an exponential growth phase. For this, nanoparticle loaded cells were washed twice with PBS (Sigma-Aldrich, Switzerland) and then detached using 450 µL Trypsin-EDTA for five minutes. Cell suspensions were transferred to tubes using 2 x 500 µL spheroid growth medium and centrifuged at 200g for five minutes. Spheroid growth medium consisted of sterile filtered cell growth medium supplemented with an additional sterile-filtered 10% FCS (total 20% FCS). After removal of the supernatant, the cell pallet was resuspended in spheroid growth medium, cells were counted and 2’500 cells per well were seeded in ultra-low attachment Biofloat 96-well plates (FaCellitate, Germany) in 100 µL spheroid growth medium in 6 technical replicates. During the initial spheroid formation and growth phase, spheroid medium was additionally supplemented with nanoparticles (TiO_2_/HfO_2_: 400 µg mL^−1^, Au100: 100 µg mL^−1^, Au50/Au25: 0 µg mL^−1^) to allow for continued loading until the day of irradiation, and to reach the desired nanomaterial uptake in spheroids that are comparable to *in vivo* and *in human* intratumoral injection masses.^42,54^ On day three after spheroid seeding, intra-spheroidal nanoparticle distribution and uptake were analyzed using SEM and ICP-OES, and spheroids were treated with or without 6 Gy of X-rays. Spheroids were irradiated with 6 Gy of 150 kVp X-rays (Seifert ISOVOLT 450, GE Sensing & Inspection Technologies GmbH, Germany) at a dose rate of ∼ 1.5 Gy min^−1^ with a phantom setup.^50^ The Control cohort received 0 Gy. Before irradiation, spheroids were washed in fresh spheroid growth medium using gentle pipetting and bright field images were taken for size assessment and normalization. Spheroids were maintained by replacing ≥ 50% of the cell culture medium every 2–3 days. For the initial dose response curve experiment (Figure 2b and c), the Au nanoparticle concentration was 50 µg mL^−1^ during the 2D cell loading and the 3D spheroid formation phases, and control or Au-treated spheroids were irradiated with 0, 2, 4, 6, 8 or 10 Gy on day 4.

### Spheroid Growth Curve Assessment

Every two to three days spheroids were imaged starting with day three after spheroid seeding. Brightfield images were acquired with a ZEISS Primovert microscope (ZEISS, Switzerland) at 4x magnification using an Axiocam 105 camera adaptor (ZEISS, Switzerland). Images were analyzed in imageJ-based open-source software Fiji (version 1.54i) using the INSIDIA macro.^55,56^ In short, the macro was used to segment the spheroid from the background and to automatically calculate its cross-sectional area at the equator. In cases where the macro was unable to accurately segment an image, cross-sectional areas were determined manually. To this end the polygon tool was to trace the edges of the spheroid in imageJ. Using the imageJ’s measure function the cross-sectional area was then calculated from the polygon selection. The Manual measurement method was checked to lead to comparable values when compared with the INSIDIA-macro. To establish growth curves, averaged cross-sectional areas from multiple spheroids per condition were plotted over time. Additionally, cross-sectional areas were normalized to initially recorded areas at day three. Cross-sectional areas were recorded until spheroids reached approximately four to five times their initial size, or until day 19 after formation, at which point the experiment was terminated. Sample size per condition was n=4 unless stated otherwise.

### Spheroid Viability

Viability of spheroids was evaluated with the CellTiter-Glo® 3D Cell Viability Assay (Promega, Switzerland). Spheroids were washed twice with 80µL of medium and transferred to a white 96-well plate (Corning, USA) in 100 µL medium. An equal amount of assay reagent was added to wells and the plate was placed on an orbital shaker set to 500 rpm for five minutes. After a further 30 minutes of incubation at room temperature in the dark, luminescence was measured with a Mithras LB 943 plate reader (Berthold Technologies GmbH & Co. KG) using 0.2 s integration time. Values were averaged from two replicates per condition and normalized to the control group.

### Spheroid Characterization

On day two and nine after formation respectively, spheroids were stained with 5 µM Hypoxia Green (Thermo Fisher Scientific Inc., USA) by removing 50 µL medium and replacing it with 50 µL of 10 µM Hypoxia Green in medium. To allow for diffusion of the hypoxia reporter throughout the spheroid, incubation took place over 24 hours at 37°C, 5% CO_2_ in a humidified incubator.^57^ Subsequently spheroids were stained using 5 µg mL^−1^ Hoechst 33342 and 50µg mL^−1^ Propidium Iodine by replacing 50 µL medium containing 10µg mL^−1^ Hoechst 33342 and 100 µg mL^−1^ Propidium Iodine. After one hour of incubation at 37°C, 5% CO_2_ spheroids were washed three times with PBS for three minutes. Following the washing steps spheroids were transferred to a glass-bottom petri dish (MatTek Corporation, USA) without prior fixation. To prevent drying out, spheroids were wetted with a few drops of PBS. Images were acquired with a Zeiss AXIO Imager M1 microscope (ZEISS, Germany). Exposure times for blue, red and green channels were set to 250 ms and kept constant between the day three and day ten spheroid.

### Spheroid Embedding and Scanning Electron Microscopy

To study the distribution of nanoparticles at the day of irradiation, spheroids were fixed at day three and imaged using scanning electron microscopy. For this purpose, cells were loaded with nanoparticles and spheroids were generated according to the described protocols. After two washes with PBS, 12 spheroids per condition were transferred to a tube. Spheroids were fixed using a solution of 2.5% Glutaraldehyde and 0.1M cacodylate buffer first for two hours at room temperature and then overnight at 4°C in the fridge. The following day, fixed spheroids were washed three times with 0.1M cacodylate buffer for three minutes. To visualize cellular structures, spheroids were stained for one hour at room temperature with 1% osmium-tetroxide in 0.1M cacodylate buffer. Thereafter three washing steps with milipore water for three minutes were performed. Spheroids were dehydrated through the following ethanol series: 30% (5 minutes), 50% (5 minutes), 70% (5 minutes), 90% (5 minutes) and 100% (3× 10 minutes). Overnight, spheroids were soaked with a 1:1 mixture of epon and 100% ethanol. Spheroids were then embedded in epon resin and transferred to a mold. Epon resin was allowed to fully polymerize in the oven at 60°C over the course of 48 hours. 200nm thick sections were obtained using a Leica EM VCG ultra-microtome (Leica, Germany). Images were acquired at 4kV using 10µs and 2x line integration on an Axia ChemiSEM Scanning Electron Microscope (Thermo Fisher Scientific Inc., USA).

### Nanoparticle Surface Functionalization for In Vivo Injections

To prepare PBS based ∼ 25 mg mL^−1^ nanoparticle suspensions injectable through 25G needles, surface functionalization had to be performed. For TiO_2_ and HfO_2_, a previously established method for the nanoparticle stabilization in complex media was used and slightly adapted.^44^ In short, ca. 100 mg of nanoparticle powder were weighted into autoclaved round bottom flasks and 20 mL of 10^−2^ *M* HNO_3_ was added. After 1 h bath sonication, 6.667 mL of a sterile-filtered 1 wt% BSA solution was added rapidly under magnetic stirring. The pH was then adjusted stepwise to neutral pH using first approx. 2 mL of 100 mM NaOH (pH ∼ 5.4) and then approx. 2.4 mL of 10 mM NaOH (pH ∼ 7.0). Thereafter, 10% of 10x PBS was added and the solutions were up concentrated using centrifugation at maximum speed (25000 g) in 2 mL tubes. Nanoparticles were finally collected into a single tube after supernatant removal and redispersed using bath sonication. The TiO_2_ and HfO_2_ nanoparticle mass concentration was quantified using ICP-OES following a H_2_SO_4_ digestion protocol described below. If necessary, the nanoparticle concentration was adjusted using 0.25 wt% BSA in PBS. The functionalization of Au nanoparticles was as following: Since citrate capped Au nanoparticles agglomerated in PBS, 25 mg mL^−1^ BSA was first dissolved in 10x PBS before 10 % of that solution was added to the 1 mg mL^−1^ aqueous Au nanoparticle stock solution. Subsequently, two centrifugation steps (6000 g, 4°C) were performed, and the supernatants were carefully taken off to reach the final up concentrated nanoparticle solution in 1 x PBS and 0.25 wt% BSA. The Au nanoparticle mass concentration was quantified using ICP-OES following an aqua regia digestion protocol described below. The final nanoparticle concentrations of TiO_2_, HfO_2_ and Au were 25.0 ± 0.7, 26.1 ± 0.4 and 24.6 ± 0.5 mg mL^−1^, respectively.

### In Vivo Experiments

1 × 10⁶ FADU cells were subcutaneously injected into the left and right flanks of female nu/nu NMRI mice (Janvier Labs, Le Genest-Saint-Isle, France). The experiment was performed according to the Ethical Committee for Animal Experimentation of KU Leuven with project number P043/2024. Mice were randomized into four treatment groups: Control (vehicle-treated), NP-treated (Au, HfO_2_, and TiO_2_), RT-treated, and a combination group (NP + RT). NPs were injected intra-tumoral at 50% tumor volume one day before the start of RT, with both the NP and control groups receiving PBS + 2.5mg mL^−1^ BSA as the vehicle solution. RT was administered once tumors reached approximately 100 mm³, with a dose of 2 Gy per fraction delivered daily for five consecutive days using the Small Animal Radiation Research Platform (SARRP, X-strahl, Camberley, UK). Dose calculations were performed using MuriPlan software (X-strahl, Camberley, UK) after cone beam CT imaging, and radiation was delivered with 220 kV photons, 13 mA, and a 10 × 10 mm or 2 cm collimator, depending on tumor size. RT response was assessed through (relative) tumor volume growth curves. Tumor volumes were measured using calipers and calculated with the formula V = (π/6) × L × W × H. Body weight and health status of the mice were monitored daily. Mice were euthanized when the average tumor volume reached approximately four times the treatment volume at start of RT or when they met the humane endpoint criteria. Maximum intensity projections of CT dicom images were performed in MATLAB (R2024b, The MathWorks, Inc., USA).

### Organ Fixation

Following nanomaterial exposure and completion of the *in vivo* experiments, the animals underwent necropsy to recover tumors along with major organs, including the liver, spleen, kidneys, and heart. The tumor tissue was fixed using paraformaldehyde (4%) for 24 h and transferred to sodium azide solution for long-term storage.

### H&E Staining and Tissue Section Imaging

The excised and 4% PFA fixed tumors were washed 3x with PBS to remove residual debris. Using a sterile scalpel, the tumor was bisected along its midline. One half was placed into a histology embedding cassette, cushioned with a soft sponge to prevent movement during the process and dehydrated using a spin tissue processor Myr STP 120 the dehydrated tumor specimen was subsequent embedded in paraffin using the tissue embedding system Shandon Histocenter 3 and solidified at 4°C. The resulting paraffin blocks were trimmed to achieve a complete tissue surface and cut into 6 µm thick fully intact cross-sections by using a microtome Leica RM2235. The tissue sections, floating in a 45°C water bath, were placed in triplicate onto Superfrost Plus Adhesion Microscope Slides. After drying, glass slides were deparaffinized for 30 min at 60°C and subsequently H&E stained using slide stainer Myreva SS-30 following standard protocols and mounted with a cover slip. Whole tumor sections were selected and 20x imaged using Slide Scanner Olympus VS200 ASW 4.1.2 using an UPLXAPO 20x/0.8 objective lens and BF default scanning mode with pre-focus in normal z plane mode for generating high resolution images captured at 0.274 µm pixel^−1^ resolution and stored as .vsi format with 24-bit RGB color depth. For scanning electron microscopy, stained sections were exposed to xylene to remove paraffin on the glass coverslip, coated with a 10 nm carbon layer (Leica EM ACE600) and imaged on an Axia Chemisem (Thermo Fisher, NL). SEM and microscopy images were overlayed using the BigWarp Plugin in ImageJ2 (Version 2.16.0/1.54p).

### Metal Quantification in Cells, Spheroids and Organs

In order to quantify nanoparticle uptake, metal mass was quantified from a cell suspension, from spheroids or from organs using ICP-OES. In case of cell monolayers, nanoparticle-loaded FaDu cells were washed using PBS, detached by 5 min of 0.5% Trypsin-EDTA treatment, and then resuspended in fresh medium after centrifugation (200g). Approximately 1 × 10^5^ cells were used per condition for analysis and transferred to quartz tubes for digestion. In case of spheroids, 6 spheroids were pooled after washing with PBS and transferred to quartz digestion tubes. In case of organs, fixed tissues were weighted, cooled to −80°C for 24 h, lyophilized and weighted, and subsequently homogenized using an achat mortar. Dry homogenized organ powder was then weighted into quartz digestion tubes. In the following, material-specific digestion protocols were used. For samples treated with HfO_2_ and TiO_2_ nanoparticles, as well as controls, digestion was performed according to a novel, HF-free, alternative digestion strategy.^58^ In brief, 1.5 mL H_2_SO_4_ was added to tubes, followed by 1 mL of H_2_O_2_ addition (stepwise addition in case of organs). During this process, tubes were immersed in a water bath to tamper the violently exothermic reaction. After 30 min, the tubes were transferred into a microwave (turboWAVE Inert, MWS GmbH) and digested at 250°C for 1h. Cell and spheroid samples treated with Au nanoparticles were fully digested in 0.1 mL H_2_O_2_ and 4 mL aqua regia (1 ml HNO_3_ and 3 ml HCL) at room temperature over three hours. Organ samples with Au were digested in the microwave after 1 ml HNO_3_ and 3 ml HCL addition and five steps of 0.1 mL H_2_O_2_ addition and mixing. All digested samples were transferred to 50 mL falcon tubes. To samples with digested gold 12.5 mL 4% L-cysteine (in 1% HNO_3_) was added coordinating gold atoms and minimizing carryover during ICP analysis. Tubes were filled to 50 mL with ultra-pure water. All samples were measured using an Agilent 5110 ICP-OES (Agilent, Switzerland) and calibrated using external standards for gold, hafnia and titanium. For calibration, standard solutions containing elements in their respective digestion matrix were used at concentrations of 0 ppm, 0.1 ppm, 0.5 ppm, 1 ppm and 5 ppm.

### Statistics

Statistical analyses of the relative tumor growth curves and Kaplan–Meier survival curves were performed using two-way ANOVA and Log-rank test. P-values of p < 0.05 were considered statistically significant.

## Acknowledgements

The authors want to thank Dr. Gabriella Éva Bodizs of the ScopeM Histology Team for her support & assistance in this work, Dr. Vera M. Kissling (Empa) for her support with osmium staining, epon embedding, spheroid sectioning and supervision of L.G.S.S. and S.H. in EM sample processing, and Dr. Davide Bottone (Empa) for his help with EM and LM image overlays. Figure 3a and 4c was partly created using Biorender.com.

S.N. is supported by a clinical research mandate from the Flemish Foundation of Scientific Research (FWO-Vlaanderen, 18B4122N).

