## Supplemental Table and Figures for "Nanoparticle Metal Mass Uptake Governs Radiosensitizing Efficacy Across 2D, 3D, and In Vivo Models"

***Table S1:*** Statistical Evaluation of spheroid dose responses corresponding to Figure 2 in the main manuscript. Spheroids were either treated with control vehicle or with Au nanoparticles. Statistical evaluation: T-Test comparing X-ray treatment conditions with the 0 Gy treatment condition within either the control (no nanoparticle) or Au spheroid group (c); ns: not significant.

| *T-Test* | | Control | | | | | Au | | | | |
|---|---|---|---|---|---|---|---|---|---|---|---|
| **Day Post Spheroid Seeding** | | **3** | **5** | **7** | **10** | **12** | **3** | **5** | **7** | **10** | **12** |
| **Day Post Irradiation** | | **-1** | **1** | **3** | **6** | **8** | **-1** | **1** | **3** | **6** | **8** |
| **P value (compared to 0 Gy)** | **4 Gy Treatment** | ns | ns | ns | * | ** | ns | ns | ** | *** | **** |
| | **6 Gy Treatment** | ns | ns | * | ** | ** | ns | ns | ** | **** | **** |
| | **8 Gy Treatment** | ns | ns | * | ** | **** | ns | ns | ** | **** | **** |
| | **10 Gy Treatment** | ns | ns | 0.13 | ** | **** | | | | | |

*ns: $p \geq 0.1$; *: $0.01 \leq p < 0.05$; **: $0.001 \leq p < 0.01$; ***: $0.0001 \leq p < 0.001$; ****: $p < 0.0001$*

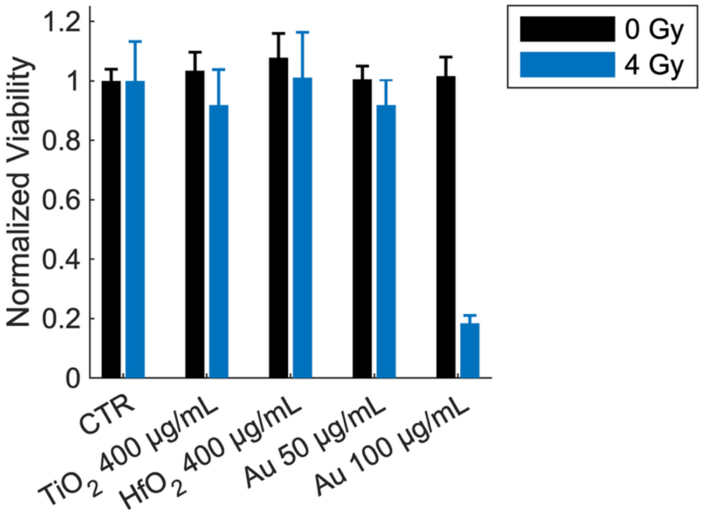


***Figure S1:*** Viability of FaDu cells pre-incubated with different nanoparticles and concentrations for 24h before irradiation with 4 Gy from a 150 kVp X-ray source. Viability was quantified 10 days post irradiation using the CellTiter-Glo® assay. Data given as Mean ± SD from two biological experiments with four technical replicates.

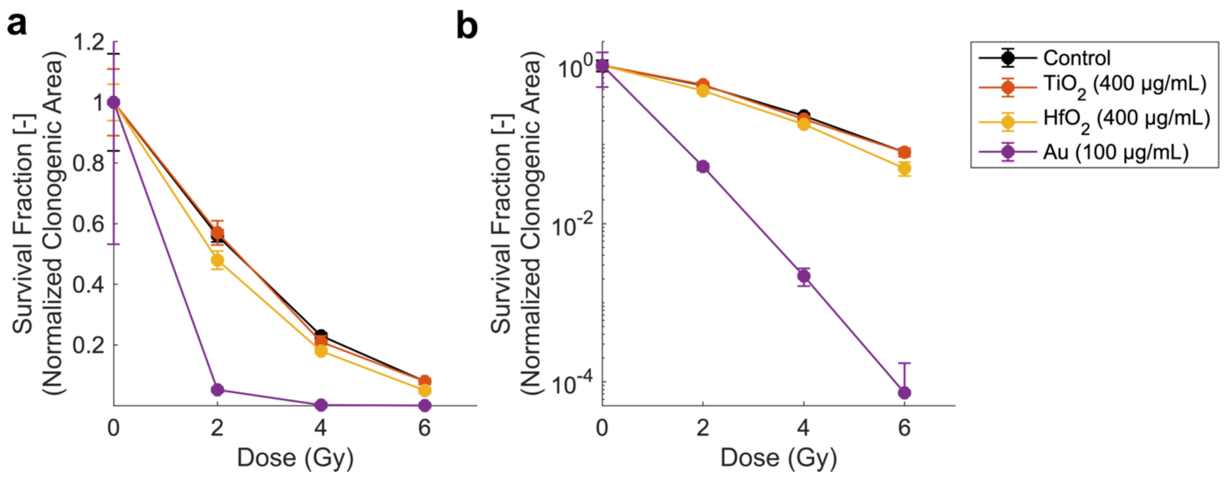


***Figure S2:*** Clonogenic Survival of FaDu cells incubated with different nanoparticles for 24h before irradiation with different Doses from a 150 kVp X-ray source. Y-axis shown in linear (a) and logarithmic (b) scale format. Data given as Mean ± SD from one biological experiment with two technical replicates.

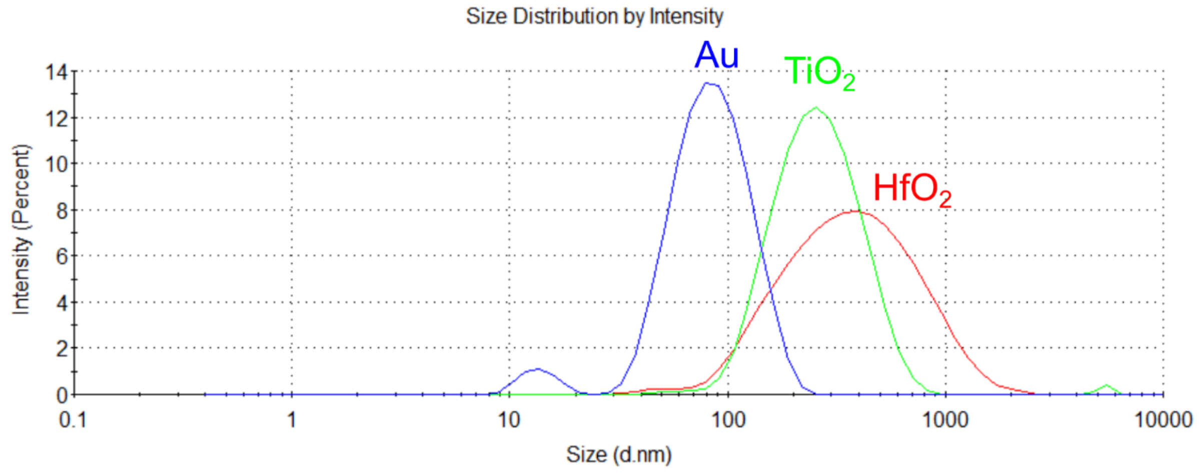


***Figure S3:*** Hydrodynamic size distribution of FCS (fetal calf serum) functionalized nanomaterials diluted in PBS measured by Dynamic Light Scattering (DLS) technique.

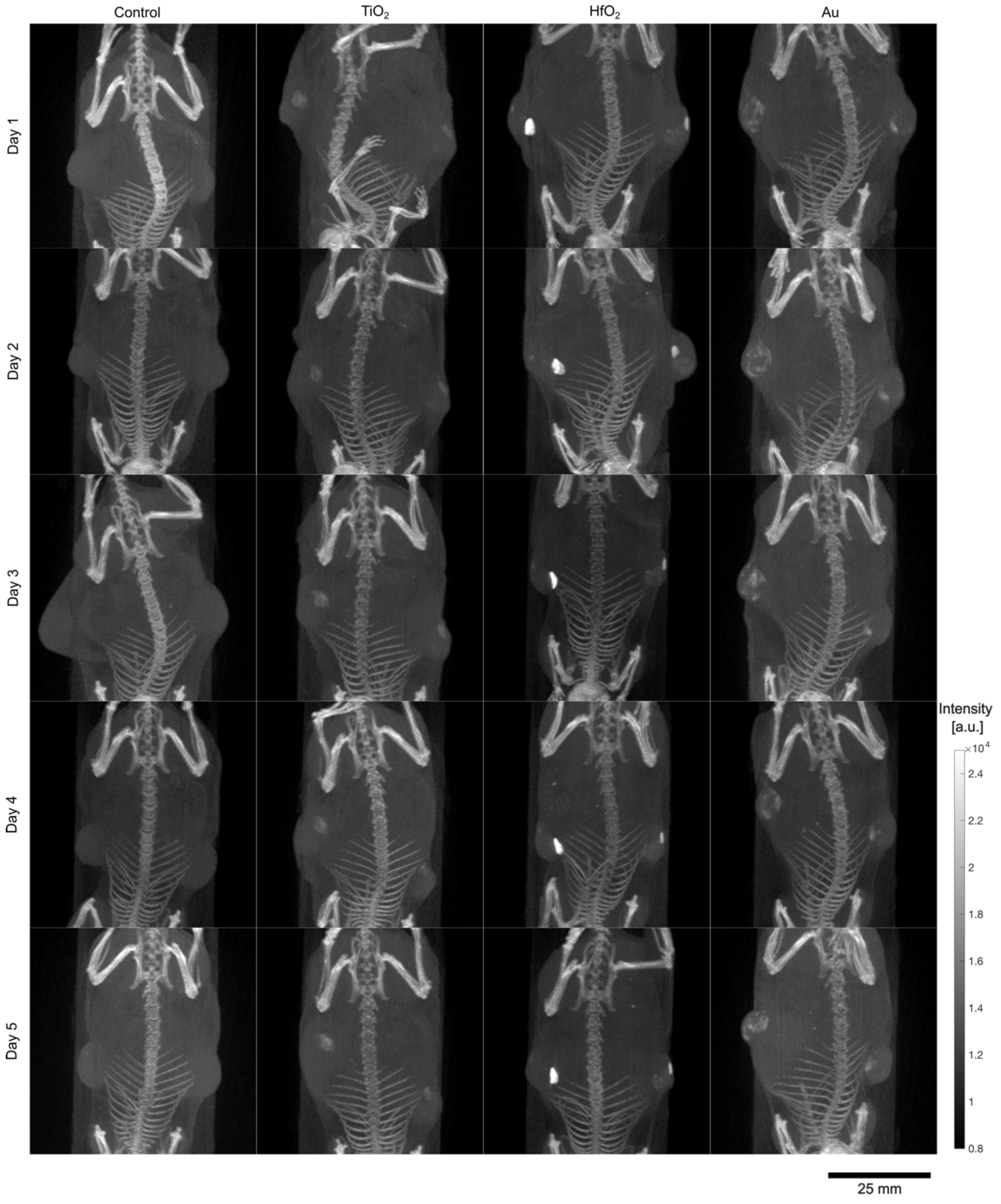


***Figure S4:*** Maximum intensity projection of CT images collected before radiation treatment on day 1, 2, 3, 4 and 5 post nanomaterial injection for each mouse (mouse 1 of 3 per nanomaterial group).

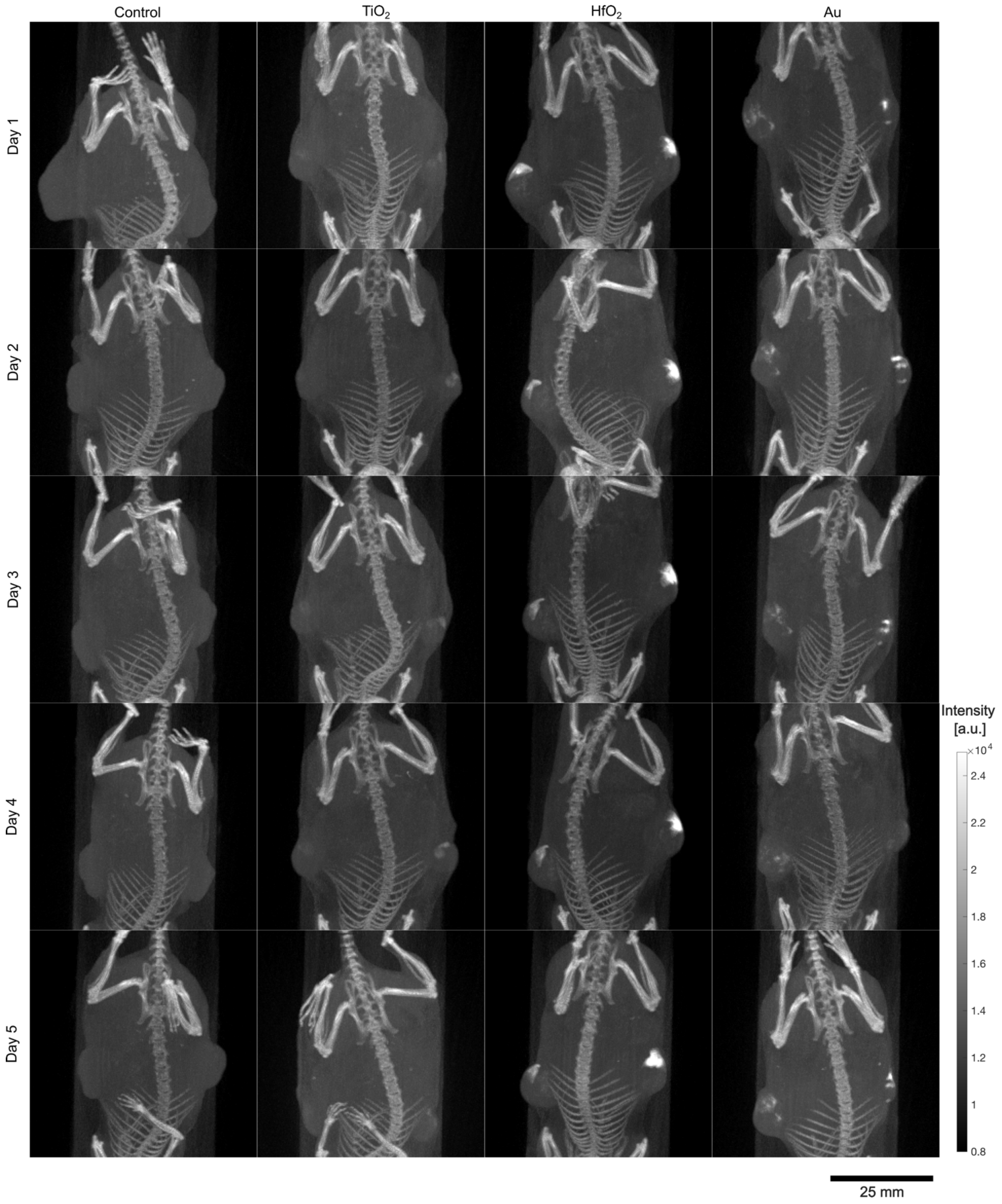


***Figure S5:*** Maximum intensity projection of CT images collected before radiation treatment on day 1, 2, 3, 4 and 5 post nanomaterial injection for each mouse (mouse 2 of 3 per nanomaterial group).

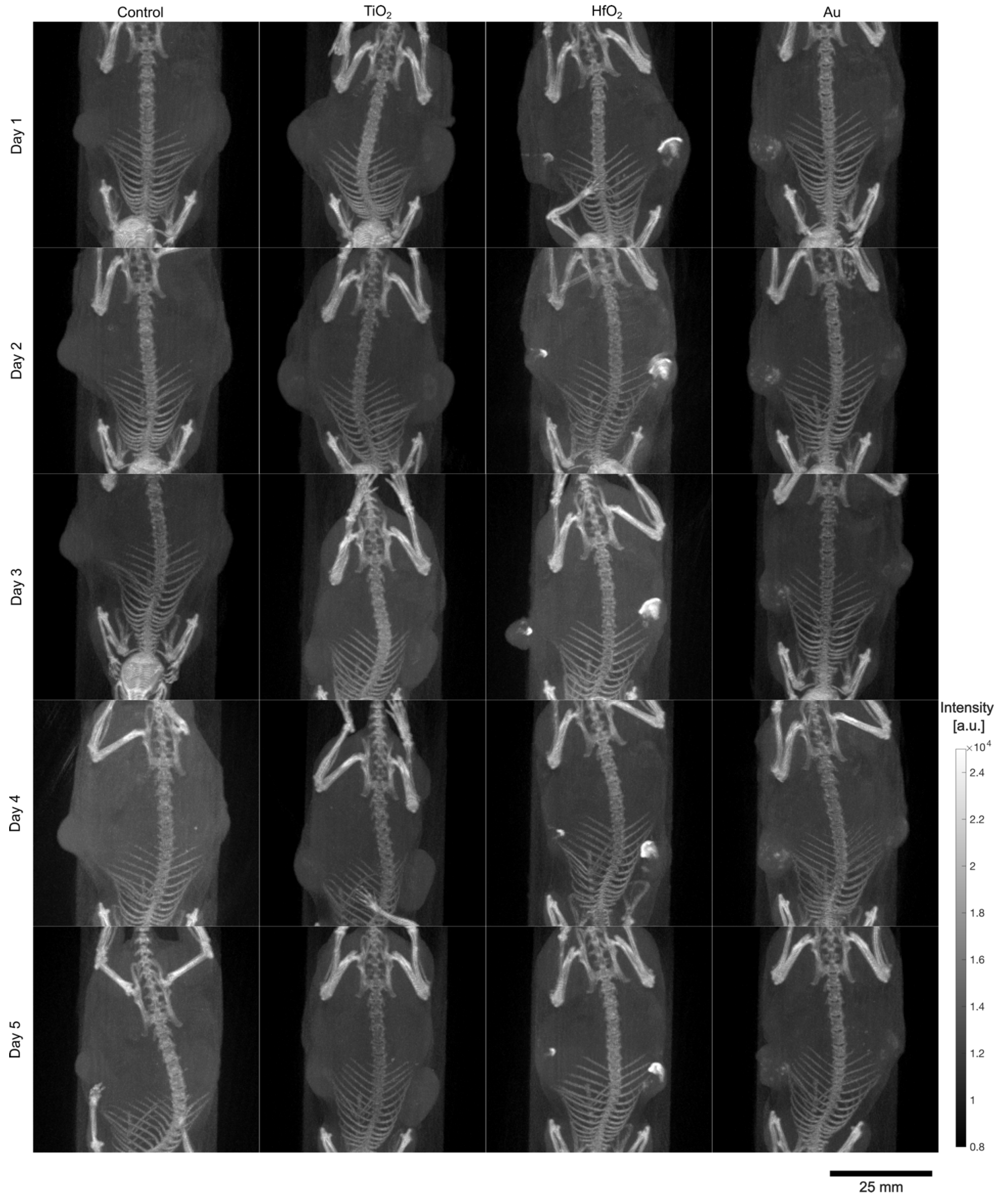


***Figure S6:*** Maximum intensity projection of CT images collected before radiation treatment on day 1, 2, 3, 4 and 5 post nanomaterial injection for each mouse (mouse 3 of 3 per nanomaterial group).
